# PBHoover and CigarRoller: a method for confident haploid variant calling on Pacific Biosciences data and its application to heterogeneous population analysis

**DOI:** 10.1101/360370

**Authors:** Sarah Ramirez-Busby, Afif Elghraoui, Yeon Bin Kim, Kellie Kim, Faramarz Valafar

**Affiliations:** Laboratory for Pathogenesis of Clinical Drug Resistance and Persistence, Biomedical Informatics Research Center, San Diego State University, San Diego, CA, USA

**Keywords:** variant calling, pacific biosciences, long reads, whole genome sequencing, mycobacterium tuberculosis, bacteria

## Abstract

**Motivation:** Single Molecule Real-Time (SMRT) sequencing has important and underutilized advantages that amplification-based platforms lack. Lack of systematic error (e.g. GC-bias), complete *de novo* assembly (including large repetitive regions) without scaffolding, can be mentioned. SMRT sequencing, however suffers from high random error rate and low sequencing depth (older chemistries). Here, we introduce PBHoover, software that uses a heuristic calling algorithm in order to make base calls with high certainty in low coverage regions. This software is also capable of mixed population detection with high sensitivity. PBHoover’s CigarRoller attachment improves sequencing depth in low-coverage regions through CIGAR-string correction.

**Results:** We tested both modules on 348 *M.tuberculosis* clinical isolates sequenced on C1 or C2 chemistries. On average, CigarRoller improved percentage of usable read count from 68.9% to 99.98% in C1 runs and from 50% to 99% in C2 runs. Using the greater depth provided by CigarRoller, PBHoover was able to make base and variant calls 99.95% concordant with Sanger calls (QV33). PBHoover also detected antibiotic-resistant subpopulations that went undetected by Sanger. Using C1 chemistry, subpopulations as small as 9% of the total colony can be detected by PBHoover. This provides the most sensitive amplification-free molecular method for heterogeneity analysis and is in line with phenotypic methods’ sensitivity. This sensitivity significantly improves with the greater depth and lower error rate of the newer chemistries.

**Availability and Implementation:** Executables are freely available under GNU GPL v3+ at http://www.gitlab.com/LPCDRP/pbhoover and http://www.gitlab.com/LPCDRP/CigarRoller. PBHoover is also available on bioconda: https://anaconda.org/bioconda/pbhoover.

## 1. BACKGROUND

The advent of whole genome sequencing (WGS) provided a long-needed research tool that is now being retooled as a diagnostic platform. On this path several challenges remain, most notably, challenges in phenotypic-genotypic associations. In infectious diseases, these challenges frequently stem from heterogeneity of the sample and from systematic errors in WGS. In this manuscript we offer PBHoover and its attachment CIGAR Roller as a consensus and heterogeneity caller which in combination with long-read sequencing offer high-fidelity heterogeneity and consensus calling without systematic bias. Amplification-based short-read WGS platforms such as Illumina dominate the current market, generally due to their lower cost. Long read sequencing, such as Pacific Biosciences’ (PacBio) Single Molecule Real-Time (SMRT) sequencing, however, has important, underutilized advantages such as absence of GC-bias and the ability to bridge repetitive elements and large structural variations (Thompson and Milos, 2011; Miyamoto *et al*., 2014; Chin *et al*., 2013; Eid *et al*., 2009). Such platforms are capable of generating long reads (2-50kb average read length) allowing *de novo* assembly and closing genomes without *a priori* structural assumptions (Ali *et al*., 2015; Yan *et al*., 2015; Steinig *et al*., 2015). Additional advantages of SMRT sequencing lie in its error profile. While these platforms suffer from higher random error rate, they do not suffer from the more difficult to correct systematic errors due to absence of the library amplification step (Loomis *et al*., 2013; Rasko *et al*., 2011; English *et al*., 2014). Random errors have a white noise profile and can be corrected by increased sequencing depth (Roberts *et al*., 2013; Koren *et al*., 2013). Systematic errors, however, are more challenging, as increased depth only increases the strength of the systematic bias.

NCBI’s Sequence Read Archive (SRA) (Leinonen *et al*., 2011) currently houses over 10,000 PacBio runs that were generated using the platform’s legacy sequencing kits (chemistry 1 [C1] and chemistry 2 [C2]). The number is significantly higher (estimated to be over 100,000 runs) when one considers all the runs that have not been uploaded to SRA. However, random errors in combination with low sequencing depth have rendered these runs unusable. Because of this, Quail *et al*., for instance, concluded that single nucleotide polymorphisms (SNP) calling using PacBio’s C1 data was inaccurate (Quail *et al*., 2012). Furthermore, using more recent PacBio chemistries, these tools allow for sensitive subpopulation detection and more accurate phenotypic-genotypic association, making the tools highly relevant to clinical applications in diagnostics and prognostics (e.g. antimicrobial drug resistance studies). In doing so, low coverage has traditionally presented significant challenges. Older PacBio chemistries, for instance, provide shallow coverage, making base and variant calling uncertain and error-prone for most reasonably-priced (e.g. using a single SMRT cell per sample for bacterial genomes) sequencing runs. Our results demonstrate that low coverage is exacerbated by aligners creating malformed CIGAR strings, causing reads to be discarded by downstream consensus callers including the manufacturer’s own software. Figure 1A demonstrates a typical variant plot across the genome of a clinical *Mycobacterium tuberculosis* (*M. tuberculosis*) isolate using PacBio’s legacy consensus calling software Evicons (Schadt *et al*., 2010; Clark *et al*., 2013; Quail *et al*., 2012)(SMRT portal version 1.3, the default tool at the time the data were produced) with manufacturer’s default settings for reference-based assembly and variant calling. This run and the subsequent analysis produced 15,689 SNPs, 261,234 insertions, and 25,834 deletions. A comparison with Sanger sequencing demonstrated that the great majority of these calls are random errors (see Results section).

**Figure 1.**
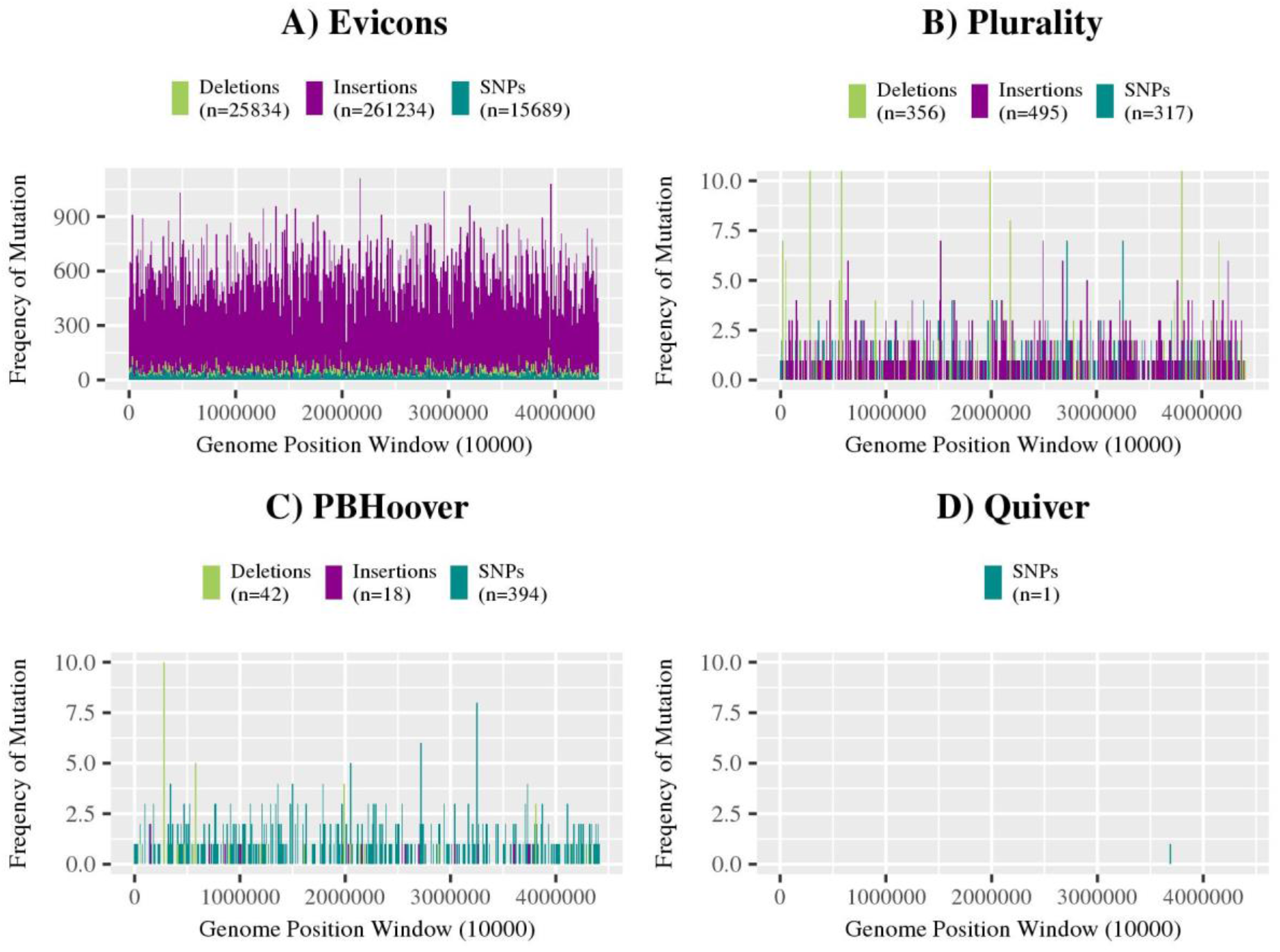
Distribution of variant calls across a clinical *Mycobacterium tuberculosis* genome (4.4 Mb) sequenced with Pacific Biosciences’ chemistry 1 on one SMRT cell called by A) PacBio’s original variant caller, EviCons; B) PacBio’s alternative variant calling algorithm, Plurality; C) PBHoover; and D) PacBio’s current variant calling algorithm, Quiver.

While this scenario can be avoided with PacBio’s more recent sequencing kits, providing greater sequencing depth, the problem reemerges in clinical applications (e.g. diagnostics, microbiome studies of infectious diseases) where sample heterogeneity splits the depth between multiple base calls. Consequently, the method presented here is developed both with PacBio’s legacy sequencing kits as well as clinical applications of the more recent kits in mind. The presented method is also readily applicable to other long-read technologies such as Oxford Nanopore. Because most currently available software is designed for consensus calling (not for heterogeneity analysis) in high-coverage, short-read scenarios (e.g. Illumina WGS), this method is also applicable with some limitations. (See Discussions).

For processing PacBio data, the manufacturer has released three consensus/variant callers, Arrow, Quiver, and Plurality (Alexander *et al*., 2011; Chin *et al*., 2013). Quiver and Arrow rely on detailed quality metrics (measured during the sequencing process) that were not recorded in older runs using legacy kits. Plurality uses a simple algorithm to call the majority “vote” of the reads at a position, but, as the authors described, (Alexander, 2016) this is an unreliable approach as it neglects total coverage and the possibility that rival calls may preclude the determination of a sensible consensus. Finally, neither of these tools were designed for heterogeneity analysis. Our tests demonstrated that none was suitable for analysis of legacy PacBio data (see Results).

To overcome these obstacles, we have developed CigarRoller and PBHoover, and optimized their use in a pipeline to call variants with high confidence on data generated from PacBio C1 and C2 and early polymerases, or from PacBio’s recent runs on clinical samples where heterogeneity may be an issue. Through comparison with Sanger sequencing, we will demonstrate our pipeline’s accuracy in consensus and heterogeneous base and variant calling with high confidence, even using C1 and C2 data, allowing the vast amount of available data to finally be accurately analyzed.

In order to demonstrate clinical relevance, in this study, we have used clinical *M. tuberculosis* strains, the causative agent of Tuberculosis (TB), the infectious disease with highest global mortality (over 1.5 million annually)(WHO, 2017). *M. tuberculosis* has a 4.4 Mb Class II genome (Koren *et al*., 2013) containing many mid-scale repeats and large repetitive regions and frequent structural variations (Mankiewicz and Liivak, 1975; Andreu and Gibert, 2008) making assembly and base/variant calling challenging. *M. tuberculosis* is known to exist in heterogeneous populations, both *in vivo* and *in vitro*—the main reason why drug resistance goes undetected and spreads (Müller *et al*., 2012). In this study, optimum pipeline thresholds have been statistically determined and utilized for detection of heterogeneous populations of *M. tuberculosis*. Several of these thresholds also agree with similar thresholds published previously (Black *et al*., 2015).

## 2. SYSTEM AND METHODS

The PBHoover pipeline is depicted in Figure 2. It includes four stages:

**Figure 2.**
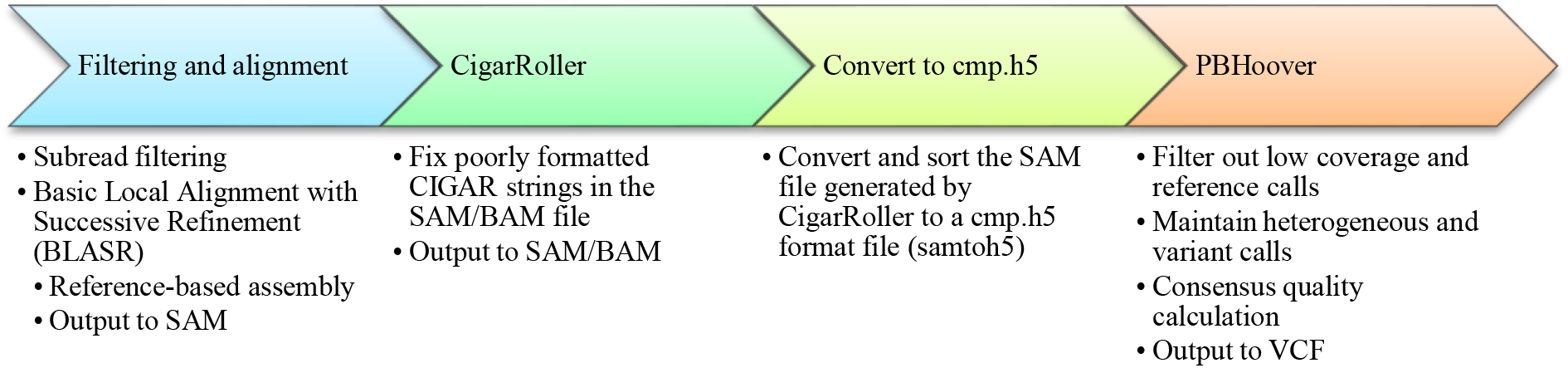
PBHoover pipeline for haploid genomes consists of four stages. The CigarRoller attachment can proceed any consensus caller independently to improve depth.

1. *filtering and alignment*: In this step, short and low-quality reads are discarded and reference-based assembly is performed by BLASR (Chaisson and Tesler, 2012).
2. *CigarRoller*: CIGAR string errors often cause problems for downstream variant callers, resulting in exclusion of the offending reads from analysis. In this step, discarded reads are examined for simple CIGAR string errors, eliminating them by merging simple cases of consecutive insertions and deletions (indels) and hence correcting creating a new local realignment, and returning the read to the pool of reads to be considered for downstream analysis (base, variant, consensus, and heterogeneity calling).
3. *conversion to h5 format*: In this step BLASR’s samtoh5 tool (Chaisson and Tesler, 2012) is used to convert the SAM file to cmp.h5 format for PBHoover.
4. *PBHoover*: uses a hypothesis-based maximum likelihood criterion to identify low coverage positions and make consensus and heterogeneous reference or variant calls.

## 3. ALGORITHM

PBHoover is a consensus and heterogeneous base and variant calling algorithm that is designed to make calls with high certainty even in most of the regions that are considered to have low coverage by other callers. As such, it is also ideal for making heterogeneous calls in regions where sufficient coverage splits across differing calls, resulting in marginal support for a consensus call and hence a “no-call” by conventional base callers. PBHoover’s performance is greatly improved by CigarRoller (see Results) and uses it to improve depth. This is consequential in regions with low coverage.

### Decision Rule for Consensus and Heterogeneity Calling

PBHoover uses a hypothesis-based Maximum Likelihood (ML) heuristic Decision Rule (DR) to make three types of consensus calls: monoclonal consensus, consensus with a minor subpopulation, and heterogeneous (multiple populations with no consensus call). The DR first develops hypotheses for each position based on aligned reads. Then, it uses a maximum likelihood model to choose the consensus call between the hypotheses. In absence of a clear “major call”, but sufficient statistical support for several hypotheses, PBHoover declares the position as heterogeneous. In presence of a major call, PBHoover still maintains a record of other hypotheses that also have sufficient statistical power. As such, PBHoover may report a major call (consensus) and a small subpopulation(s) for some positions.

In this manuscript, the term “heterogeneous” refers to a stricter meaning. Namely, we use the term only for positions that have sufficient statistical power for several hypotheses, but lack a clear major call. For such positions, all subpopulations are reported but no consensus call is made. In case of insufficient support for any hypothesis, the position is labeled as “low-coverage.” As such, it is entirely possible that a position with marginally sufficient coverage be labeled as low coverage by PBHoover if the reads split across multiple calls.

### Hypothesis-based maximum likelihood model

An important distinction between PBHoover’s decision rule and that of other callers is in its consideration of mixed populations. PBHoover considers multiple alleles at each position and hence does not dismiss reads that disagree with the most prevalent call as sequencing error. Rather, it examines the hypothesis that they may be representing distinct subpopulations. This has important implications in probability models and QV score calculations since disagreeing reads at a position may no longer reduce the certainty of the major call (variant or reference) and, if they meet the Minimum Certainty Requirement (MCR) (see next section), but rather contribute towards the certainty of a subpopulation. For each position of the reference sequence, PBHoover considers six possible hypotheses for six possible outcomes: A, T, C, G, insertion, and deletion. If more than one hypothesis proves to be statistically significant (meets MCR), the position will be labeled as indicative of mixed populations. If on the other hand, only one of the six hypotheses meet the MCR, the position will be labeled as “monoclonal.”

### Minimum Certainty Requirement (MCR)

The MCR is the minimum number of supporting reads that a hypothesis needs for it not to be dismissed as sequencing error. It is based on the error profile of the library chemistry and a user provided parameter, *maximum allowable consensus error rate* (MACER). Because of differing likelihoods of error for different types of variants, the statistical model provides stricter MCR for types with higher likelihood of error (single-base indels [SBI] for PacBio chemistries).

Because MACER indirectly defines what should be considered as low coverage, it also indirectly defines the percentage of the genome with marginal coverage. As such, this parameter can also be used to estimate sequencing yield (after application of all certainty filters) based on genome size, number of SMRT cells, and chemistry/polymerase error profile. In our experiments we have used a single SMRT cell for *M. tuberculosis* genome (4.4Mb) using C1-C1, early C2 (C2-C2), and P4-C2 polymerase-chemistries on RS and RSII PacBio instruments.

In this study, we offer a default MACER value of one error in 100 base calls (QV20) for bacterial genomes. This is in-line with criteria suggested in the literature (Underwood and Green, 2011). It is important to recognize that the method described here assumes the absence of systematic error. See “Applicability to other platforms” in Discussions for application of PBHoover to data generated by sequencers with sequencing bias such as Illumina.

EQ1 estimates the probability that the reads supporting a hypothesis are due to sequencing error.

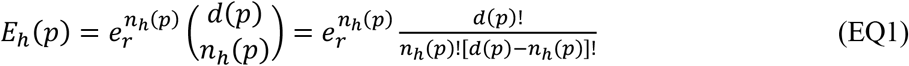

We denote *n_h_*(*p*) to be the number of reads supporting hypothesis *h* at position *p* of the reference genome. As EQ1 describes, by assuming a random error profile, this probability is simply the raw read error rate, *e_r_*, to the power of number of reads supporting the hypothesis multiplied by the binomial distribution of *n_h_*(*p*) over *d*(*p*) (total sequencing depth at position *p*).

For C1, *e_r_* is reported to be as high as 17.9% (Ferrarini *et al*., 2013; Chin *et al*., 2011), while C2’s rate has been reported to be as high as 15% (with polymerase 4) (Frank *et al*., 2016), and ~11% was reported for later polymerase-chemistry combinations (e.g. P6-C4) (Korlach, 2015). The Phred scale quality score for hypothesis *h*, can then be estimated by the standard calculation as shown in EQ2 (DePristo *et al*., 2011):

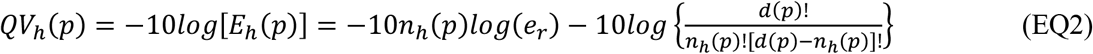

It is important to recognize that EQ2 estimates hypothesis-specific quality values, distinct from consensus quality scores which will be estimated later in this article. EQ2 can be used to numerically estimate MCR for various error types by replacing *E_h_*(*p*) with the MACER rate specified by the user and replacing *e_r_* with the raw error rate for each type of error. Because SBI errors are about four times more likely in C1 chemistry (Table 1 and Figure 1), and overall raw C1 read error rate is estimated to be 17.9%, the *e_r_* of SBI is estimated to be 14.3% and that of SNPs and multi-base indels to be 3.6%. Figure 3 depicts the relationship between hypothesis QV, sequencing depth, and MCR for C1 Chemistry. As it can be seen, for SNPs and multi-base indels (Figure 3A), to stay above the MACER value of QV20 in regions of the genome that have a total depth between four and 12, at least three reads (MCR value) need to support a hypothesis. At depth of 13, three reads are no longer sufficient, as the QV score dips below the specified MACER value of QV20. Therefore, regions with depths between 13 and 22 require at least four reads, and depths greater than 23 (but less than 32—not shown in the Figure), require at least five. This trend of increasing MCR with increasing depth continues for higher depths. As Figure 3B depicts, for SBIs, due to higher probability of such error, EQ2 results in stricter MCR values. For depths between four and 12, six reads are required, while depths between 13 and 15 require seven reads supporting a hypothesis. Again, the trend continues for SBIs at stricter levels as depicted by Figure 3B. For more recent chemistries (not shown in the figures), lower raw read error rates result in lower MCR. As such, greater sensitivity in detecting heterogeneity can be offered for these chemistries.

**Figure 3.**
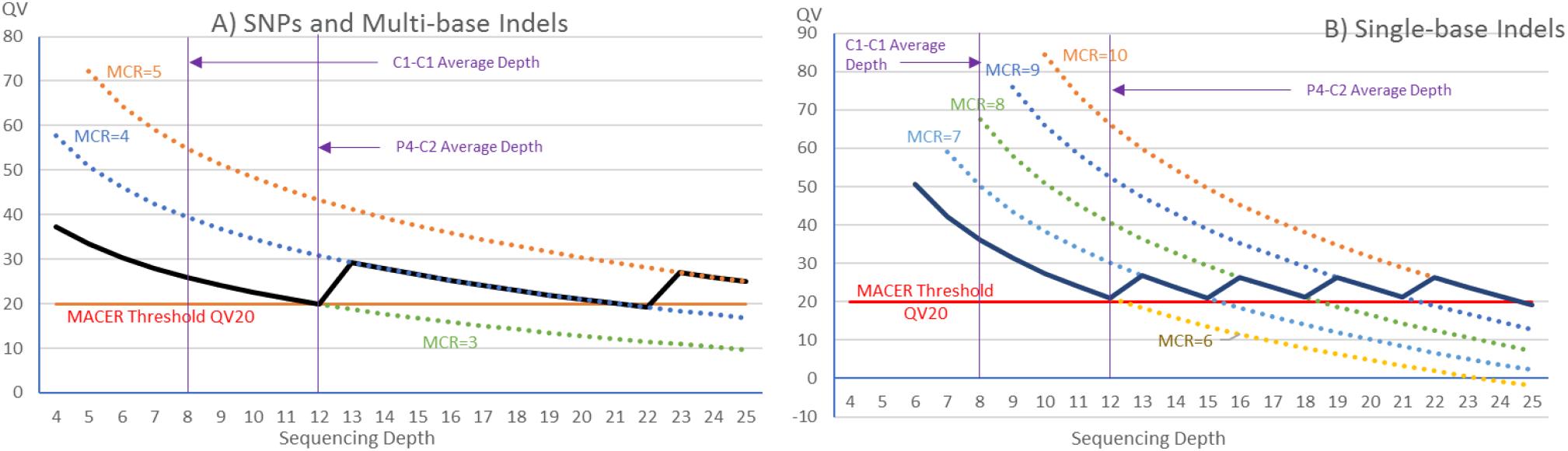
Relationship between MCR, hypothesis QV, and sequencing depth of C1 Chemistry for *M. tuberculosis* (4.4 Mb). One SMRT cell is used per sample with a genome-wide average depth of 8X. Average depth of 12X for P4-C2 is also shown (second vertical line) for comparison. Raw sequence error rate of 14.3% was used for single-based indels (B). 3.6% was used for all other types (A). Solid curves indicate the MCR used by PBHoover for single-base indel calls (B) and other types (A) for sequencing depths up to 25X. MRC requirements would be lower (not shown) for C2 chemistry due to lower error rates. MCR curves are calculated based on C1 error rates.

**Table 1.** Median variant counts per variant caller across 348 *M. tuberculosis* clinical strains sequenced on Pacific Biosciences RS platform using chemistry 1 (C1) or chemistry 2 (C2). One isolate was run once on C1 and once on C2. For this isolate the reads from both runs were combined (C1+C2 column). Only Plurality and PBHoover were tested on this isolate. “-“ indicates that the method was not tested for the category.

| Variant Caller | SNPs |  |  | Insertions |  |  | Deletions |  |  |
| --- | --- | --- | --- | --- | --- | --- | --- | --- | --- |
|  | C1 | C2 | C1+C2 | C1 | C2 | C1+C2 | C1 | C2 | C1+C2 |
| **EviCons** | 6,071 | 5,372 | - | 56,436 | 53,263 | - | 16,776 | 13,154 | - |
| **Quiver** | - | 774 | - | - | 65 | - | - | 42 | - |
| **Plurality** | 802 | 1,188 | 1441 | 329 | 111 | 136 | 1,696 | 1,646 | 1,914 |
| **PBHoover** | 843 | 1,194 | 1426 | 32 | 67 | 103 | 334 | 376 | 485 |

### Consensus calling and heterogeneity detection

Figure 4 depicts PBHoover’s DR for consensus and heterogeneity calling. Table 2 displays examples of how DR works. In the context of consensus calling, we distinguish between monoclonal calls (Examples 1 and 2 in Table 2), major calls with minor subpopulations (Examples 3 and 4), and heterogeneous calls where no clear major call exists (Example 5). Anytime a consensus call is made (variant or reference), it is reported according to the DR (Figure 4) along with any subpopulations that meet their MCR (not shown in Figure 4). If no subpopulation is reported, that consensus call is considered monoclonal.

**Figure 4.**
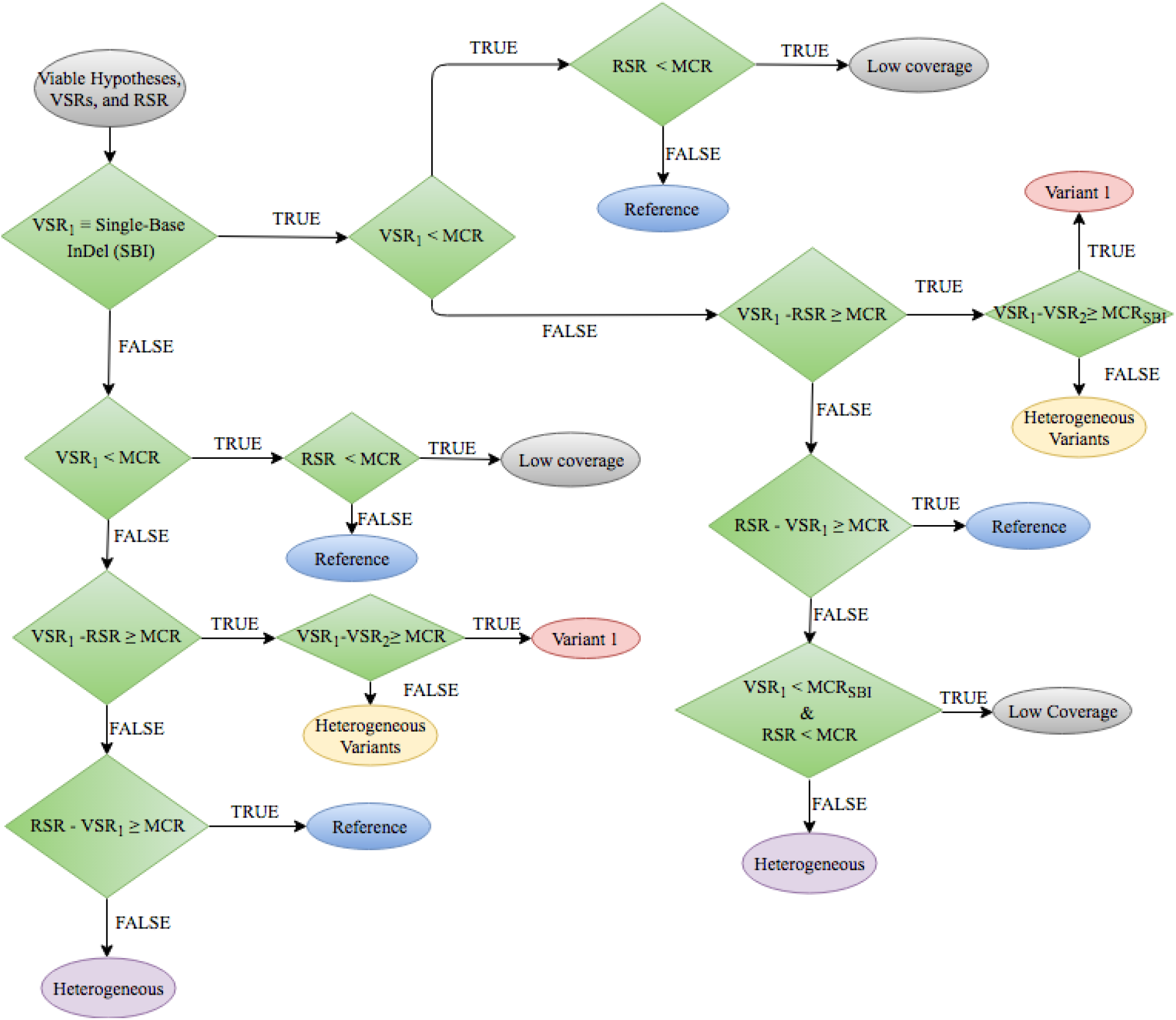
PBHoover consensus calling criteria for consensus calling. RSR: reference supporting reads. VSR: variant supporting reads. VSR_1_: Number of reads supporting the most prevalent variant. VSR_2_: Number of reads supporting the second most prevalent variant. MCR: Minimum Certainty Requirement for SNPs and multi-base indels. MCR_SBI_: MCR for SBIs. Viable hypotheses are those that have equal to or greater than MCR support. “Variant 1” indicates a consensus call for the most prevalent variant. “Heterogeneous” call indicates sufficient and similar support for the reference as well as a variant. “Heterogeneous Variants” indicates that the top two hypotheses belong to two different variants and that there is similar level of support for the two. VSR and RSR of nonviable hypotheses are set to zero for any downstream calculations as the reads supporting those hypotheses are dismissed as sequencing error.

**Table 2.** Examples of PBHoover consensus calling. A is the reference call, MCR=4 and MCR_SBI_=6 for all Examples. H1: Hypothesis 1; H2: Hypothesis 2; H3: Hypothesis 3. Notation: A:11 indicates 11 reads call for A at the position; RSR: Reference Supporting Reads; VSR: Variant Supporting Reads.

| Example | H1 (REF) | H2 | H3 | RSR | VSR<sub>1</sub> | VSR<sub>2</sub> | Consensus Call | Consensus Status |
| --- | --- | --- | --- | --- | --- | --- | --- | --- |
| **1** | A:7 | G:3 | C:2 | 7 | 0 | 0 | A | Monoclonal: Reference A |
| **2** | A:2 | G:52 | C:2 | 0 | 52 | 0 | G | Monoclonal: Variant G |
| **3** | A:7 | G:52 | C:2 | 7 | 52 | 0 | Ga | Variant G; wildtype (a) subpopulation |
| **4** | A:11 | G:4 | C:2 | 11 | 4 | 0 | Ag | Reference A; mutant (g) subpopulation |
| **5** | A:7 | G:10 | C:5 | 7 | 10 | 5 | gac | Heterogeneous: g, a, c |
| **6** | A:3 | G:3 | C:2 | 0 | 0 | 0 |  | Low Coverage |
| **7** | A:7 | G:4 | Δ:6 | 6 | 6 | 4 | agΔ | Heterogeneous: a, g, deletion |

As mentioned before, if no hypothesis meets MCR, the position is labeled as “low coverage” (Example 6). The distinction between a consensus call with a subpopulation and a heterogeneous call is in the presence of a clear major population (Example 3) or lack thereof (Examples 5 and 7).

As Figure 4 depicts, if *VSR*_1_ − *RSR* ≥ *MCR* the consensus call will agree with the reference and if *RSR* − *VSR*_1_ > *MCR*, PBHoover will call for a variant. If *VSR*_1_ ≥ *MCR* and, |*RSR* − *VSR*_1_| < *MCR* or *VSR*_1_ − *VSR*_2_ < *MCR*, the consensus call will not be possible and the position will be called heterogeneous (Examples 5 and 7, Table 2). Figure 4 shows two types of heterogeneity conditions which could have diagnostic significance: one is where there is sufficient support for the reference as well as a variant, and the second is where there is not enough support for the reference but there is sufficient support for two different types of variants. Here, it is important to keep in mind that before the evaluation of these conditions, RSR and VSR values are set to zero, if they do not meet their respective MCRs. *Consensus quality score*: Since the consensus is called based on all hypotheses that meet MCR, the quality score will depend on the possibility that any of the hypotheses are incorrect. As such, an upper bound for the consensus error at position *p* can be estimated by EQ3:

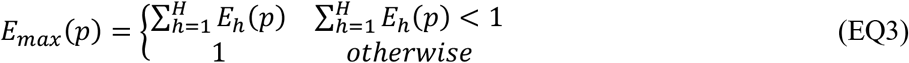

This is an upper bound because the probability of error of various hypotheses are not mutually exclusive and hence they could add up to be more than one. For this reason, EQ3 caps the sum at one. The lower bound for the consensus quality score is calculated by EQ4:

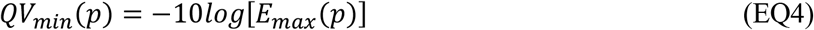

The *QV_min_* of multi-base indels is calculated by taking the lowest *QV_min_* of the positions involved in the variant after merging the indels.

### Merging Variants and Partial Local Realignment

PBHoover’s analysis of the genome, one position at a time, could lead to non-contextual variant calling. For instance, SNPs could be presented as a deletion and a subsequent insertion, or a deletion of multiple bases could be presented as multiple consecutive single-base deletions. As a remedy for this issue, we have implemented a post-processing step to fix this and several other common issues, so to accurately represent variants. The goal of this step is to combine variants that appear in consecutive positions into one variant. This step only performs partial realignment to address this issue for specific combinations of variants and not for all possible combinations. Supplementary Table 1 lists the combinations that are addressed by the realigner.

To address the issue of improving the alignment of reads in the presence of deletions and insertions, the partial realigner perform a sequence of steps as follows: First, indels are left-aligned using bcftools norm (Li *et al*., 2009); second, consecutive SNPs are combined to create a multi-base substitution record (e.g. Supplementary Table 1, Case 1). Next, consecutive deletions are combined into one multi-based deletion event (Supplementary Table 1, Case 2). The fourth step merges deletion-SNP events (Supplementary Table 1, Case 3). The fourth step attempts to combine consecutive insertion-deletion and deletion-insertion events (Supplementary Table 1, Cases 4 and 5). The consecutive records are then merged only if the length of the insertion is equal to the length of the deletion. The resulting single or multi-base substitution is placed at the position of the first single base variant that was merged. Finally, three specific patterns of three variants that are frequently observed are merged. These are: INS-SNP-DEL, DEL-SNP-INS, and SNP-INS-SNP (e.g. Supplementary Table 1, Cases 6-8).

Because these steps could create additional opportunity for merging variants, these three steps are repeated until the alignment converges and no more instances of the addressed types are detected.

## 4. IMPLEMENTATION

The implementation details of the three preprocessing steps depicted in Figure 2 is described below:

1. *Read filtering and Reference-Based Assembly*: Raw reads were filtered using manufacturer’s recommended thresholds (minimum read score of 0.75, minimum subread length of 50bp, minimum read length of 50bp), then mapped to *M. tuberculosis* H37Rv (Genbank accession NC_000962.3) for all clinical samples. Ambiguously mapped reads (mapping quality value of less than 20) were excluded from the assembly. Manufacturer-recommended reference-based mapping software BLASR v2.0.0 (Chaisson and Tesler, 2012) was used to filter and then align the remaining reads to the corresponding reference strain. The resulting alignment of each read is coded as a “CIGAR” string and stored in BAM format. Unconventional formatting of CIGAR strings often causes problems for downstream variant callers resulting in discarding reads with such problems.
2. *CigarRoller*: CigarRoller was created to fix poorly formatted CIGAR strings produced by BLASR (Chaisson and Tesler, 2012). Variant callers such as GATK require properly formatted CIGAR strings according to a specific set of rules and discard nonconforming reads (McKenna *et al*., 2010). Five types of formatting issues have been observed in CIGAR strings: differing alignment and cigar length, hard/soft clips in the middle of the CIGAR string, reads starting or ending with deletions, fully hard or soft clipped alignments, or consecutive indels (McKenna *et al*., 2010). CIGAR strings generated by BLASR frequently have the consecutive indel issue (e.g. 1D1I, 2D3I1D). Because C1 and older C2 data already have low depth of coverage, the discarding of reads due to poorly formatted CIGAR strings decreases coverage to a point where a consensus cannot be called. CigarRoller, currently, only fixes CIGAR strings that contain up to three consecutive indels and discards reads with more than three consecutive indels. Hard or soft clips and mismatching lengths were never observed in our data. CigarRoller is developed as an independent module and can be used independently for other purposes as well. CigarRoller can accept either SAM or BAM files as input and can produce output in either format. For two consecutive indels, a shorter indel is combined with a longer indel, and the combined matching position is added to the neighboring number of matches. For example, the CIGAR string 3D2I1M (three deletions, two insertions, and one mismatch) becomes 1D3M. In instances of three consecutive indels, merging from left or right creates different results (e.g. 1I2D2I, from the left: 2M1I; from the right: 1I2M). Since neither CigarRoller nor PBHoover perform true local realignment, CigarRoller alternates between left merge and right merge to reduce bias.
3. *Conversion to H5 format*: PBHoover requires an H5 formatted file as input to retrieve all quality scores, including base, insertion, and deletion quality values. BLASR’s samtoh5 (Chaisson and Tesler, 2012) module was used for conversion of BAM to cmp.h5.

### Verification of PBHoover’s accuracy

C1 (n=75) and C2 (n=274) runs of 348 (one was sequenced on both chemistries) clinical strains of *M. tuberculosis* (isolated from patients in India, Moldova, Philippines, and South Africa) were sequenced on PacBio RS and RSII instruments and were used to test CigarRoller and PBHoover.

Genome Analysis Toolkit (GATK) was used to test CigarRoller’s performance. Percentage of reads discarded by GATK before and after CigarRoller was compared to assess effectiveness of the module. For PBHoover, we first compared its performance to three other manufacturer developed variants callers, Evicons (Schadt *et al*., 2010; Clark *et al*., 2013; Quail *et al*., 2012), Quiver, and plurality (Alexander *et al*., 2011; Chin *et al*., 2013). Next, Sanger sequencing was used as the “gold standard” to evaluate the accuracy of PBHoover calls. Targeted regions of all isolates’ genome were subject to Sanger sequencing (Rodwell *et al*., 2014). These regions were chosen because of their (genes *inhA* promoter, *katG*, *ahpC*, *rpoB*, *rrs*, *eis* promoter, *tlyA*, *gyrA*, and *gyrB*) involvement in antibiotic drug resistance for clinical relevance. Supplementary Table 2 lists the coordinates of the sequenced regions in the reference genome. In total 866,477 bases were sequenced by Sanger across the 348 isolates.

For PacBio runs, we used one SMRT cell for all but for 24 isolates where we used more than one SMRT cell for experimentation. In such cases all reads from the multiple SMRT cells were combined into one alignment run. We determined the number of concordant base calls between Sanger and PBHoover, which includes matching variant and reference calls. To quantify the discordance, we determined the number of false positive (variants call by PBHoover but reference by Sanger) and false negative (variant call by Sanger but reference by PBHoover) calls. We excluded positions of low coverage from this analysis.

## 5. RESULTS

### CigarRoller

On average, across 75 C1 runs, 68.9% of the reads were used by GATK and 31.1% were discarded. CigarRoller salvaged 99.6% of these (31% of all reads) and discarded only 0.35% of them (0.11% of all reads). This represents a 45% improvement (from 68.9% to 99.89%) in sequencing yield in terms of usable read count. Across the 274 C2 runs, on average, GATK discarded nearly 50% of all reads. CigarRoller salvaged 98% of the discarded reads and only discarded 0.28% of all reads. This represents a nearly 100% improvement in read count yield. The salvaged reads all contained two or three consecutive indels and discarded reads all harbored more than three consecutive indels. The other types of CIGAR string issues were never observed in 348 runs.

### PBHoover

#### Comparison to other variant callers

A single C1 run (Figure 1) demonstrated why Evicons and Quiver were no longer considered for analysis of C1 data. The number and types of variant calls for three PacBio-developed variant callers in the single run is displayed in Figure 1. Plurality was the only algorithm with results within the expected range (Figure 1B). Evicons (deprecated) (Figure 1A) and Quiver (Figure 1D) called an unrealistic number of variants, 302,757 and only one, respectively.

As the single run predicted, the three manufacturer callers produced vastly different results on the rest of the isolates. To assess the use of PBHoover on PacBio C1 data, we compared variants called by all four callers across all isolates. Table 1 represents median counts per algorithm per variant type for all four callers across the 348 isolates. EviCons noticeably overcalls all variants for C1 data. Quiver is not optimized for C1, therefore, only C2 data was analyzed. Due to the low coverage areas in C2 data, Quiver was not able to call as many variants as PBHoover. Plurality calls more indels than SNPs. The majority of these are single-base sequencing errors, which Plurality is not sensitive to. Previous studies have shown that number of SNPs in a typical clinical *M. tuberculosis* genome far exceeds number of indels. (Wang and Chen, 2013; Merker *et al*., 2013)

#### Variant quality scores

Figure 5A presents the consensus QV scores for each type of variant called. The variant graph demonstrates similar QV scores at lower depths but a clear bifurcation between indels and SNPs at higher depths, with indels exhibiting higher QV scores than SNPs.

**Figure 5.**
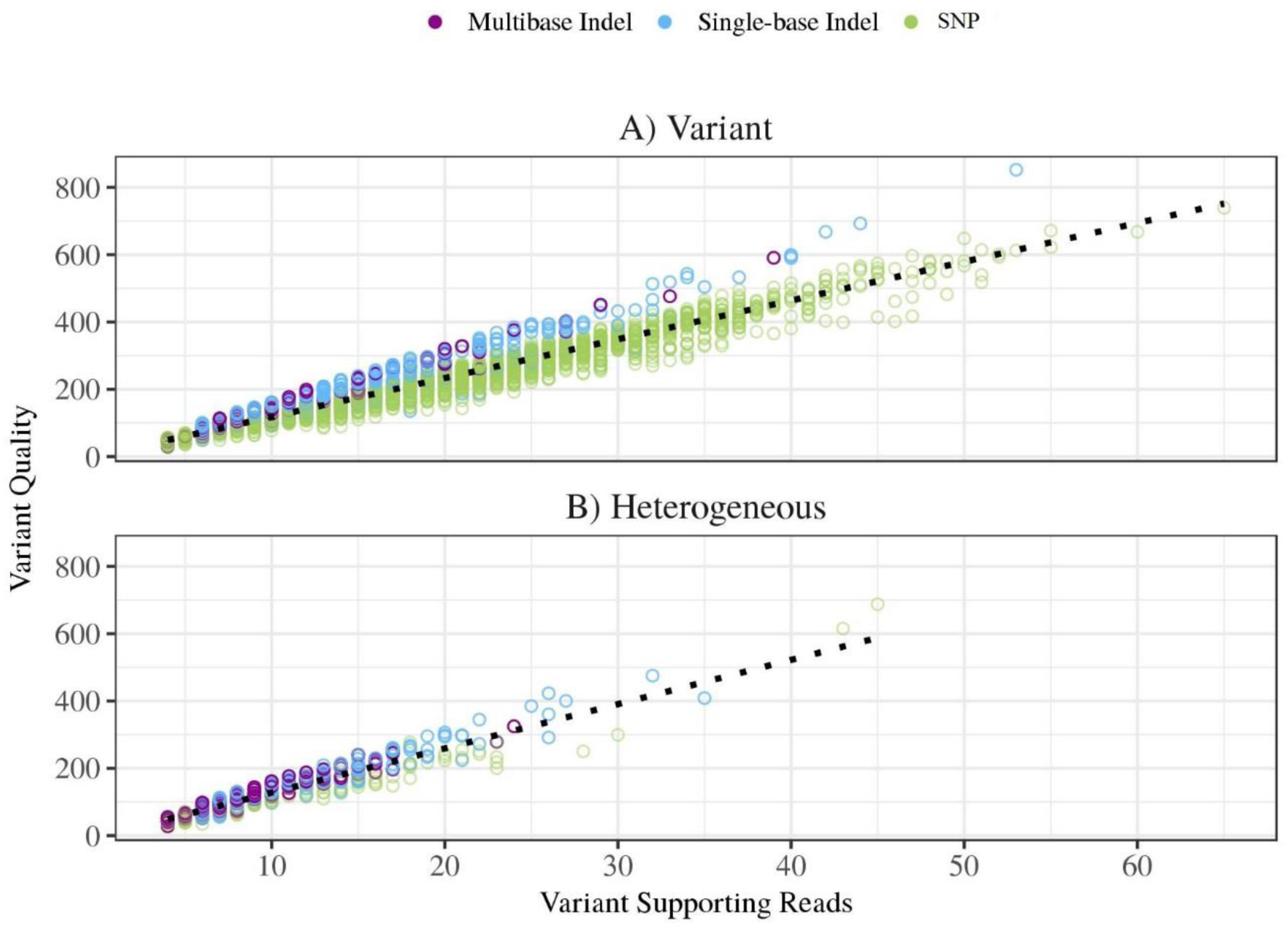
Consensus variant quality in relation to variant supporting reads for the A) positions where a major variant could be called and B) heterogeneous calls where a major variant could not be identified, made by PBHoover in a C2 run of a *M. tuberculosis* clinical isolate.

Figure 5B demonstrates an example of the amount of heterogeneous calls and corresponding VSR and consensus QV. The pattern seems to be similar to that of consensus calls in Figure 5A with the difference in that heterogeneous calls are far fewer and have overall lower QVs. This is an expected pattern. It is important to note that the figure only shows the consensus QV which is based on the QV of all subpopulations (EQ4). The figure does not display the number of subpopulations.

#### Consensus Accuracy

To assess the consensus accuracy of PBHoover, we used Sanger Sequencing as the gold standard. Sanger sequencing results for the clinical isolates included in this study was published previous (Rodwell *et al*., 2014). Sanger sequencing was only performed for part of each genomes. The regions sequenced by Sanger is listed in Supplementary Table 2. We compared all PBHoover consensus calls within these regions to those of Sanger calls. Figure 6 displays the results of this comparison.

**Figure 6.**
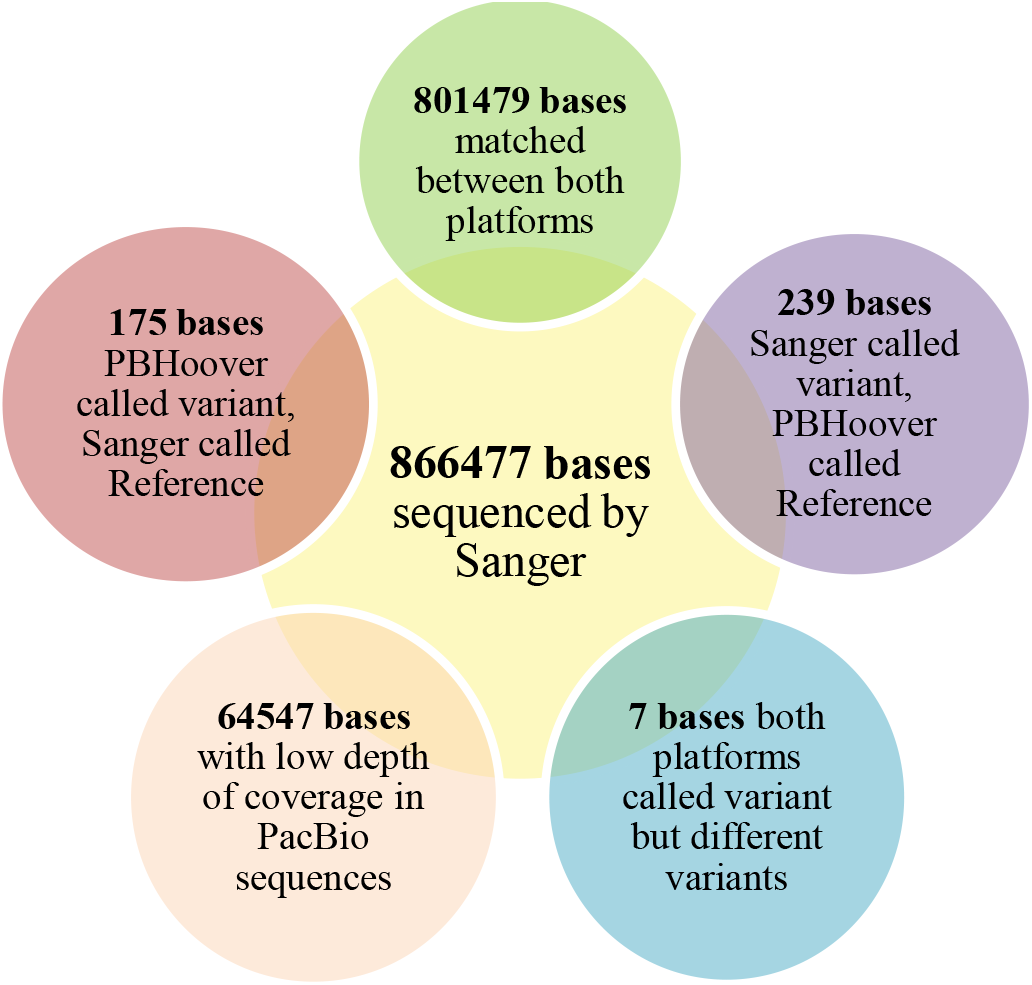
Comparison of Sanger sequencing and PBHoover calls from Pacific Bioscience’s chemistries 1 and 2 sequencing on *Mycobacterium tuberculosis* clinical isolates (n = 348). Low depth positions were excluded from this analysis.

In summary, 99.95% (QV33) of PBHoover calls were concordant with those of Sanger (Figure 6). In total, across all 348 isolates, 64,547 positions were labeled as low coverage by PBHoover and therefore were excluded from accuracy analysis. Of the remaining 801,930 consensus calls, 175 were false positives and 239 were false negatives. The number of false negatives were expected to be higher than false positives since we stay well above the MACER value of QV20 for most positions (Figure 3) and therefore implementing stricter MCR for all calls. In seven cases both methods called a variant, but they called different variants. This is likely due to heterogeneity. As such, the total error count was 421 out of 801,930 consensus calls, resulting in an average error rate of 0.05% (QV33), well above industry standards.

#### Heterogeneity

Among the bases sequenced by sanger, 30 positions were identified as heterogeneous by PBHoover. Reference was called by Sanger for 22 of these positions, while seven had a concordant variant call (Sanger variant call agreed with one of the subpopulations detected by PBHoover), and one position had a discordant variant (no subpopulation detected by PBHoover had the Sanger variant).

## 6. DISCUSSION

Reliability and sensitivity of alignment, base, variant, and consensus calling are important factors in the accuracy of WGS-based diagnostics. Heterogeneity, a natural phenomenon in pathogens, complicates the genotypic-phenotypic relationship and hence reduces the sensitivity of molecular diagnostics. Ignoring heterogeneity could lead to incorrect diagnosis, weaker genotypic-phenotypic relationships, and subsequent misled conclusions such as existence of alternative mechanisms of disease (or drug resistance), when in fact the real cause lies in a small subpopulation with the common mechanism. PBHoover and its attachment, CigarRoller, are designed to address the reliability and sensitivity issues. These qualities make the PBHoover suite the preferred tool for consensus, base, variant, and heterogeneity calling on noisy and/or low depth sequencing data (e.g. PacBio’s C1 and C2 runs). For high fidelity and/or high depth data (e.g. the more recent PacBio chemistries) the suite outperforms other callers where heterogeneity is a possibility.

For those who have turned away from low-coverage or noisy sequencing data (Quail *et al*., 2012), PBHoover offers a way to salvage and analyze their data. For those who are in search of alternative molecular mechanisms, PBHoover offers a more careful look at whether there is sufficient genotypic-phenotypic evidence for the existence of such mechanisms through heterogeneity analysis.

CigarRoller has demonstrated its ability to salvage reads to a great degree (31% of C1 and 49% of C2 reads), only discarding a small percentage of reads (0.11% in C1 and 0.28% in C2). The salvaged reads would have been discarded by most callers (e.g. GATK). CigarRoller is required for PBHoover, but also can be used in other sequencing pipelines as a general-purpose alignment clean-up tool.

### Consensus accuracy

PBHoover base calls closely matched Sanger sequencing (QV33), with rare exceptions. Here, we assumed Sanger as the gold standard and the truth. However, Sanger sequencing is known to suffer from amplification bias in the PCR step and polymerase slippage in tandem repeats and homopolymers (Kircher and Kelso, 2010). Consequently, some of the discordance may be either due to PBHoover’s ability to observe heterogeneity or Sanger sequencing error. Nevertheless, assuming PBHoover error in all discordant cases, results in 99.95% concordance with Sanger, providing important support for PBHoover.

### Heterogeneity detection

An important aspect of PBHoover is the ability to detect subpopulations. This is especially important with pathogens as the subpopulation may have gained the ability to subvert specific pressures, such as the immune system or antibiotics/antivirals. Once the specific pressure is applied (e.g. treatment commences), this population is selected for and becomes the dominant strain causing the infection. Detecting these subpopulations early in infection and treatment will assist with future treatment regimens, promote positive patient outcomes, and help prevent outbreaks. As the comparison with Sanger demonstrated, 73% (22 of 30) of the heterogeneous cases are called reference by Sanger. This is a major issue in diagnosis, since for example in the case of drug resistance, the 22 would be classified as wild type (drug susceptible), resulting in standard treatment, giving the resistant subpopulation time to infect others.

### Comparison to other callers

EviCons was the original variant caller for PacBio data and demonstrated a high rate of variability. EviCons can analyze and call variants on low coverage data, but the thresholds to call a variant were extremely loose or lacking (Table 1 and Figure 1).

Although Quiver is the current and recommended algorithm for local realignment and consensus and variant calling for PacBio data, it significantly underperforms with C1 data (Figure 1), and was not designed to detect subpopulations. Quiver’s inability to call variants in C1 data is likely due to low coverage depth in legacy data and the lack of quality values introduced in more recent sequencing software versions.

Plurality is a simple but strict “majority vote” algorithm, considering the call with the highest number of supporting reads, similar to PBHoover but without consideration of the platform error profile or heterogeneity. Because of the high rate of SBIs in PacBio data, Plurality makes more erroneous indel calls. This is clearly exhibited in Figure 1B and Table 1.

### Sequencing yield and depth

For C1-C1 runs (75 genomes), which had the lowest coverage, using one SMRT cell per sample, on average, had the following yield after all quality filters and CigarRoller improvements: 40% (1.8Mb) of the genome had sufficient coverage (9X or better), 52% (2.3Mb) had marginal coverage (between 4X and 8X), and 8% had low coverage (3X or less). As low as these percentages seem, through comparison with Sanger sequencing, we demonstrated that PBHoover can still make accurate calls even in most of the marginal coverage regions.

The percentage of the genome that falls into the sufficient, marginal, and low coverage regions change depending on the stability of the genome, heterogeneity of the population, and chemistry. As Figure 3 and the data presented in this manuscript demonstrate, if PBHoover is used for consensus and variant calling, one SMRT cell per sample is sufficient for *M. tuberculosis* even using C1 Chemistry. In order to use PBHoover effectively for prokaryotes with larger genome sizes, we recommend the number of SMRT cells to be determined in a way that each run produces at a minimum 8X average coverage. For smaller genome sizes or when using the newer Sequel platform, multiplexing provides a cost-effective solution.

### Applicability to more recent PacBio data

More recent PacBio chemistries have lower error rates and greater depth. As such, PBHoover’s ability to make base calls in low coverage regions becomes less relevant but offers far greater sensitivity in detecting mixed populations. EQ2 can be used to estimate this sensitivity. Assuming MACER value of QV20, depth of 80, and raw read error rate of 11% for P6-C4 (Korlach, 2015), PBHoover’s MCR for SNPs will be seven reads (QV21). This leads to the discovery of a subpopulation that is only 9% of the total population (assuming uniform sampling). At the depth of 150 (average depth for P6-C4 for our experiments is 120), MCR will be 11 and subpopulation detection will be at 7%. This is in line with phenotypic techniques’ sensitivity to detect heteroresistance to most drugs (Cambau *et al*., 2015) and better than some (e.g. MGIT’s sensitivity of 10% in detecting heteroresistance to pyrazinamide (Glader *et al*., 2015)) in tuberculosis, making amplification-free genotypic detection of heteroresistance a viable diagnostic alternative. Finally, for recent chemistries, the assumption of higher prevalence of SBI errors (ratio of 4-to-1 with respect to other errors) needs to be revisited for more accurate modeling.

### Applicability to other platforms

PBHoover’s stochastic models assume random error events. Systematic errors violate this assumption. As such, the application of PBHoover to platforms such as Illumina where amplification bias could result in systematic error, would remove random errors, but not systematic errors. Thus, in this sense, the expected performance of PBHoover in consensus calling will be in par with other callers. For heterogeneity detection, and the consequential genotypic-phenotypic associations, PBHoover is expected to outperform other callers. To use PBHoover with other platforms’ data, raw read error rate, and the error profile (i.e. relative frequency of each error type) need to be adjusted to model the new platform accurately.

### Limitations and other considerations

An important aspect that is lacking in the PBHoover pipeline is complete local realignment. Quiver retains an algorithm for local realignment, which may alter specific calls due to misalignments. Although PBHoover captures subpopulations with great sensitivity, this sensitivity needs to be improved. The solution to this is higher depth which is a reality with PacBio’s more recent chemistries. This offers an attractive solution for amplification-free heterogeneity analysis.

## 7. CONCLUSIONS

With whole-genome sequencing reaching popularity, analyses and pipelines to discern significant biological consequences are of utmost importance. Longer read platforms have the unique ability to capture repetitive regions and to close genomes, a more recent accomplishment. However, PacBio’s legacy data presented several obstacles that we have been able to overcome, although low depth will always be a challenge. With CigarRoller and PBHoover, we have shown that we are able to retrieve much more information (including heterogeneity) from older chemistries and polymerases than PacBio’s variant caller, Quiver (Chin *et al*., 2013), and obtain high concordance with data from Sanger sequencing. For newer chemistries, PBHoover’s heterogeneity detection is a strong advantage bringing sensitivity of molecular heterogeneity detection in par with (and in some cases better than) phenotypic detection methods.

## 8. COMPETING INTERESTS

Authors have no competing interests. The funding bodies had no role in the design of the study or collection, analysis, and interpretation of data or in writing the manuscript.

## 9. AUTHOR’S CONTRIBUTIONS

SRB implemented the PBHoover algorithm, debugged all codes written for this project, helped in developing a test suite for the code, packaged PBHoover and its attachments for public availability, and took the lead in writing the manuscript. AE helped in the development of the algorithm, implementation of it, writing a test suite for the code, and helped in packaging of the code for public availability. YBK designed, implemented, and wrote a test suite for CIGAR Roller. KK designed, implemented, and wrote a test suite for partial local alignment. FV designed the algorithm for PBHoover and helped SRB in writing the manuscript.

## 10. ACKNOWLEDGEMENTS

This work was funded by a grant from National Institute of Allergy and Infectious Diseases (NIAID Grant No. (R01AI105185). All authors were supported by this grant. SRB and A.E. were also supported by through scholarships from a National Science Foundation Grant (no. 0966391).

